# In Silico Evaluation of Biomarker Genes for Melanoma Carcinoma

**DOI:** 10.1101/2023.01.29.526121

**Authors:** Bharat Kwatra, Mohammed Moin Khan

## Abstract

Melanoma skin cancer is a primary cutaneous malignancy. Melanoma cancer of the skin cell is one of the major skin cancer worldwide, ranking first in frequency. Melanoma skin cancer has been strongly associated with psoriasis. Here, we use computational methods in an effort to identify possible biomarkers for psoriasis related Melanoma skin cancer. To do this, we downloaded gene expression microarray data from the GEO (Gene Expression Omnibus) database in the GSE series: GSE14905 and pre-processed it in the Bioconductor repository for R. The data was screened for DEGs using a rigorous methodology, which included the use of statistical testing methodologies and tools (Differentially Expressed Genes). Psoriasis has 6749 up-regulated genes and 7142 down-regulated genes. Psoriasis DEGs were combined with the NCG dataset resulting 874 up-regulated genes and 74 down-regulated genes for an in depth analysis of how differential expression can lead to malignancy. we used stringDB diseases dataset in Cytoscape to generate network of melanoma proteins, which were further mapped to DEGs and igraph to construct a GRN. In addition, module level analysis was carried out because of the benefits it provides in terms of stability and comprehension of intricate GRNs, resulting 17 biomarkers (17 up-regulated). There is an emphasis on the network’s topology as well. The findings suggest that the network has a hierarchical structure. Additionally, survival analysis of 8 biomarkers results obtained from intersection mapping of skin cancer stringDB and biomarkers, was carried out. Significant enrichment of KEGG pathways was found. They also illuminate the interplay between biomarkers whose upregulation may lead to melanoma skin cancer. These findings may inform future investigations and the identification of potential therapeutic targets for this condition as well as shows the potential link between psoriasis and melanoma.

## 1. INTRODUCTION

Malignant melanocytes (melanoma carcinoma) is a primary malignancy of melanocytes (skin pigment). It is a rare malignancy with a dismal outlook that affects just 22 out of every 100,000 Americans (Cancer statistics from the Centre for Disease Control and Prevention). Though it only makes up 4% of skin cancer incidence, it is responsible for 75% of skin cancer fatalities [1]. Melanocytic nevi and ultraviolet radiation seem to be the main reasons for this malignancy [2]. More people are being diagnosed with melanoma than any other kind of cancer. While the exact percentage growth from year to year varies by population, it has been anywhere between 3% and 7% [3]. Breslow thickness, the frequency of ulceration and regression, and the percentage of male patients are all poor prognostic indicators, and they increase with age. However, as people become older, the risk of developing a metastatic SLN decreases. This may reflect a distinct biological behaviour (hematogenous spread) of melanomas in older patients, or it may indicate that the SLN method has a lower sensitivity (greater false-negative rate) in this population [4] [4]. Melanoma has been linked to psoriasis more than any other skin condition, according to recent research [5]. Surveillance Epidemiology and End Results (SEER) data shows that melanomas account for a disproportionate share of deaths from skin cancer in the United States. Analysis of biological data via a computational lens may provide light on the nature of the illness. Differential gene expression between psoriasis and melanoma has been shown via comparative research. In this study, we analyse data from gene expression microarrays in R, stingDB, and Cytoscape to construct a Gene Regulatory Network. In addition, we identify subnetworks and communities, and then follow the network to track down the biomarkers. We also do analyses of survival, as well as analyses of cluster enrichment and GO enrichment.

## 2. METHODOLOGY

### 2.1. Pre-processing of Data

The microarray information was obtained from NCBI’s GEO (Gene expression Omnibus, http://www.ncbi.nlm.nih.gov/geo/) database. Using the term ‘psoriasis’ in conjunction with the matching GSE series, we did a step-by-step search to identify human gene expression patterns associated with psoriasis. Datasets that included a comparison of normal and control tissues were favoured (GSE14905) [6]. The GEOquery tool in bioconductor was used to pre-process and download microarray data [7]. R packages from Bioconductor were imported into the environment for grouping data, normalising results, and eliminating noise. Later expression set was generated for further analysis.

### 2.2. Screening Differentially Expressed Genes

DEGs were extracted from the gene expression matrix. After distinguishing infected samples from control samples, we averaged gene expression levels for each probe number. After a platform GPL570-based gene-probe correlation was established, probe numbers from the expression profile could be transformed into their corresponding gene symbols. Using the VOOM and LIMMA pipelines [8], we found the fold change (FC) by subtracting the mean values of infected samples from those of healthy controls. The cut-off point was set at 0.50. Additionally, DEGs were derived by filtering FC values Network of Cancer Genes (NCG) was then used to check them for cancer causing genes (Network of Cancer Genes, ncg.kcl.ac.uk)[9].

In addition, a list of genes linked to melanoma cancer was compiled by doing a Cytoscape [10]. string disease query for the term “melanoma”, resulting 2000 genes, The final list of DEGs was obtained by intersecting NCG filtered DEGs and melanoma genes.

### 2.3. Network Construction and Topological Properties

Bioconductor’s stringDB package [11] and igraph [12] were used to map the intersected cancer genes discovered by NCG (Network of Cancer Genes) and the Melanoma gene query onto a gene regulatory network. In addition, Cytoscape was used to create a network of these genes, which is a popular use of the programme for mapping attribute data onto a protein-protein interaction network or metabolic pathway, for example. The STRING database programme (included in Cystoscope’s public database area) was used to build the network. In addition, we took into account the GRN’s topological properties by drawing plots for the degree distribution (node-degree distribution P(K)), the clustering coefficient C(K), and the neighbourhood connectivity CN(K), as well as the centralities (betweenness CB(K), closeness CC(K), and eccentricity CE) (K) for all 948 NCG obtained genes.

### 2.4. Finding Modules

Importantly, the GRN was then used to produce possible modules. We used the R and iGraph [12] package to create the modules, the cluster package to create the clusters, and the cluster edge betweenness technique to determine the modularity and transitivity of the resulting graphs. In order to locate modules, a dendrogram was constructed from these groups. By following a set of instructions in R, we were also able to produce text files containing the gene names of the individual modules. Biomarkers were identified in the form of genes in these modules.

### 2.5. Gene Enrichment, KEGG Pathway and Survival Analysis

#### 2.5.1. StringDB enrichment

StringDB enrichment analysis was performed in order to obtain the function of gene network [12].

#### 2.5.2. Gene Ontology enrichment

Gene Ontology enrichment, for Biological process, Molecular Function and Cellular Component was done using R and cluster profiler package [12]

#### 2.5.3. KEGG pathway search

KEGG pathway search was done was performed using DAVID 6.8 (Database for Annotation Visualization and Integrated Discovery, https://david.ncifcrf.gov/) [13].

#### 2.5.4. Survival Analysis

cBioPortal was used to preform survival analysis of each biomarker(8) hub on skin cutaneous melanoma Pan Cancer Atlas [14, 15]

## 3. RESULTS AND DISCUSSIONS

### 3.1. Differentially Expressed Genes

Geo dataset contained 82 samples for psoriasis, out of which 21 samples were normal and 43 samples were diseased. Figure 1 shows the normalized sample boxplots, further dimensionality reduction was done shown in Figure 2 UMAP plot. VOOM results for precision weights and mean variance trend weights group is shown by Figure 3. Upregulation and downregulation of proteins is shown Figure 4. Total 6749 up-regulated genes and 7142 down-regulated genes were found.

**FIGURE 1.**
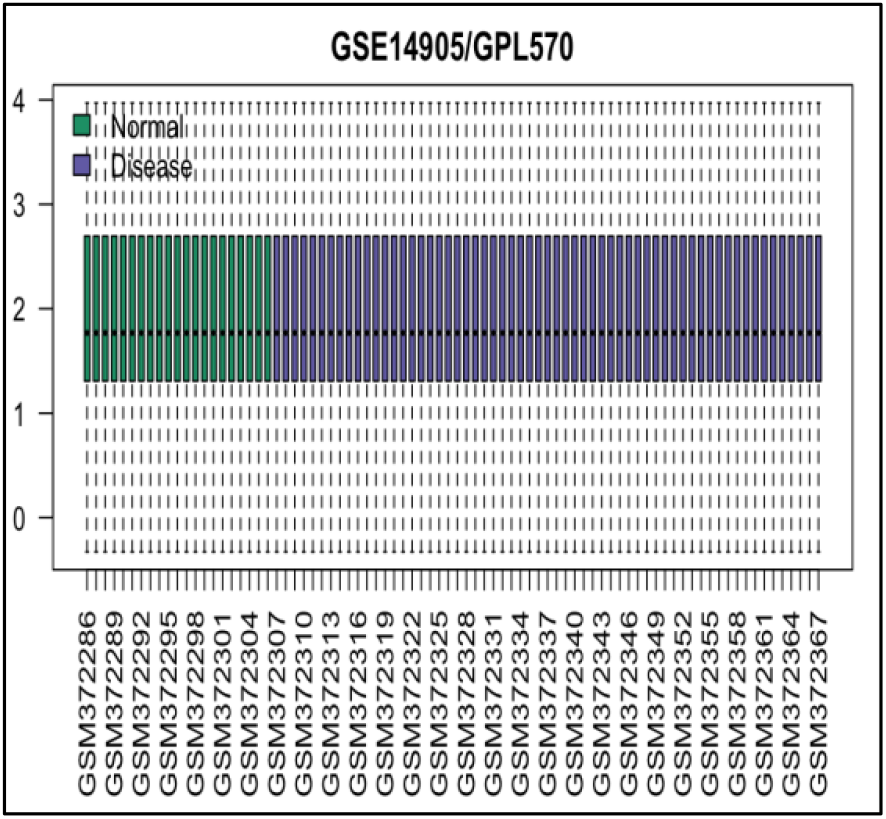
SHOWS NORMALISED SAMPLE DATA

**FIGURE 2.**
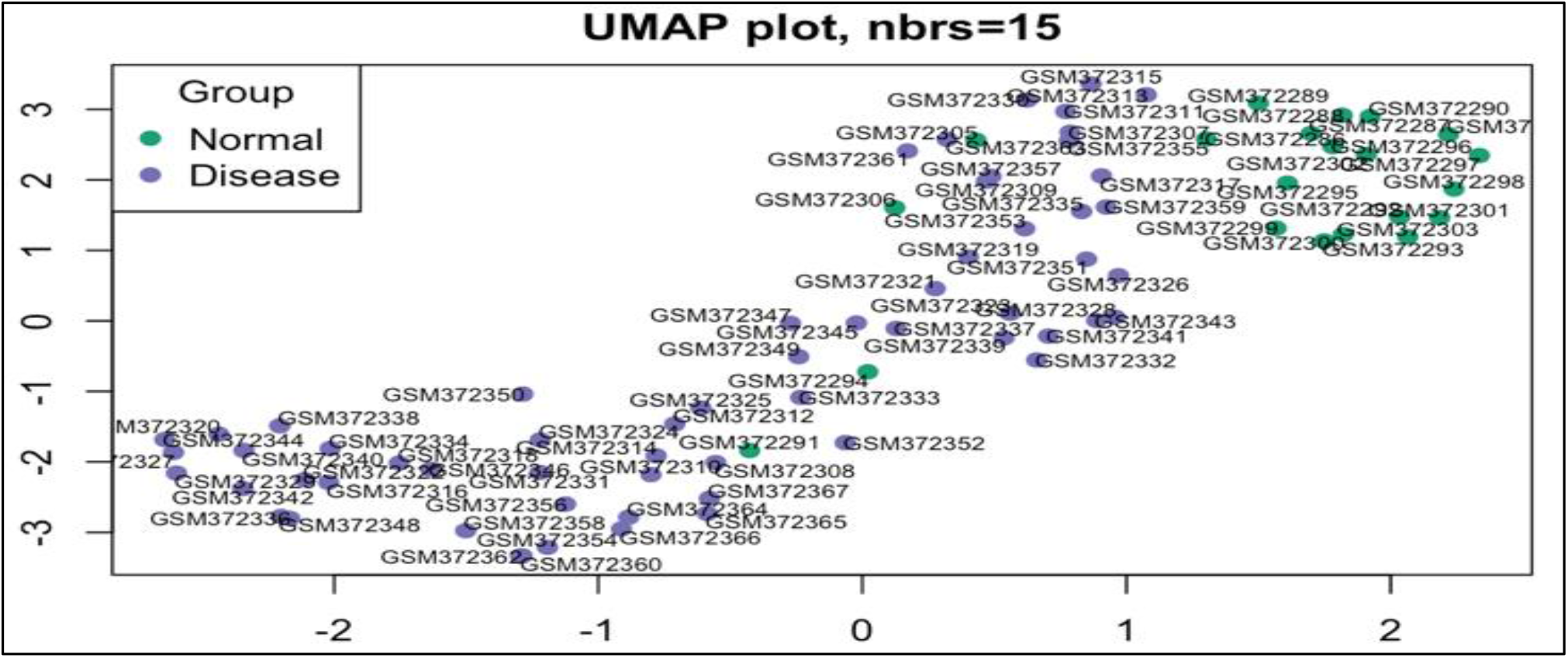
SHOWS UMAP PLOT OF SAMPLE DATA

**FIGURE 3.**
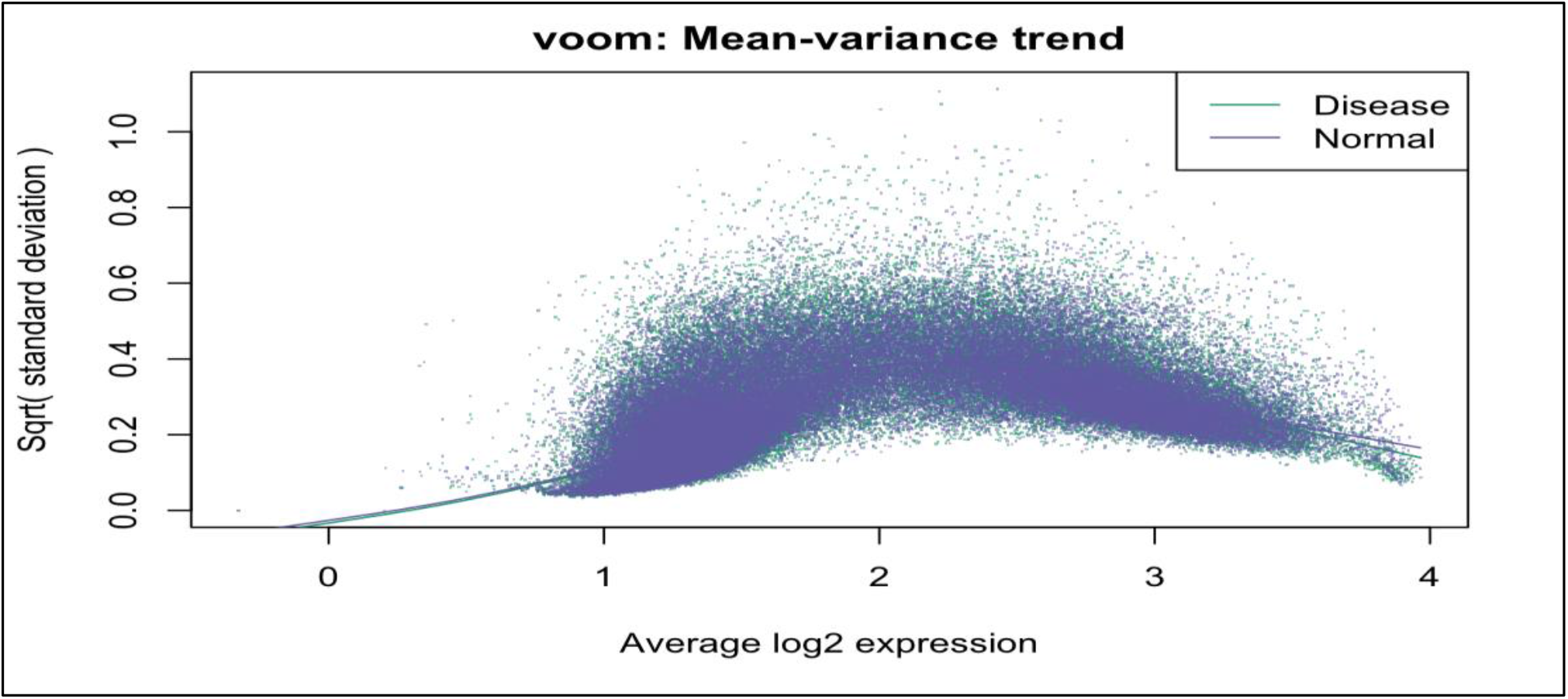
SHOWS VOOM WEIGHTS BY GROUP

**FIGURE 4.**
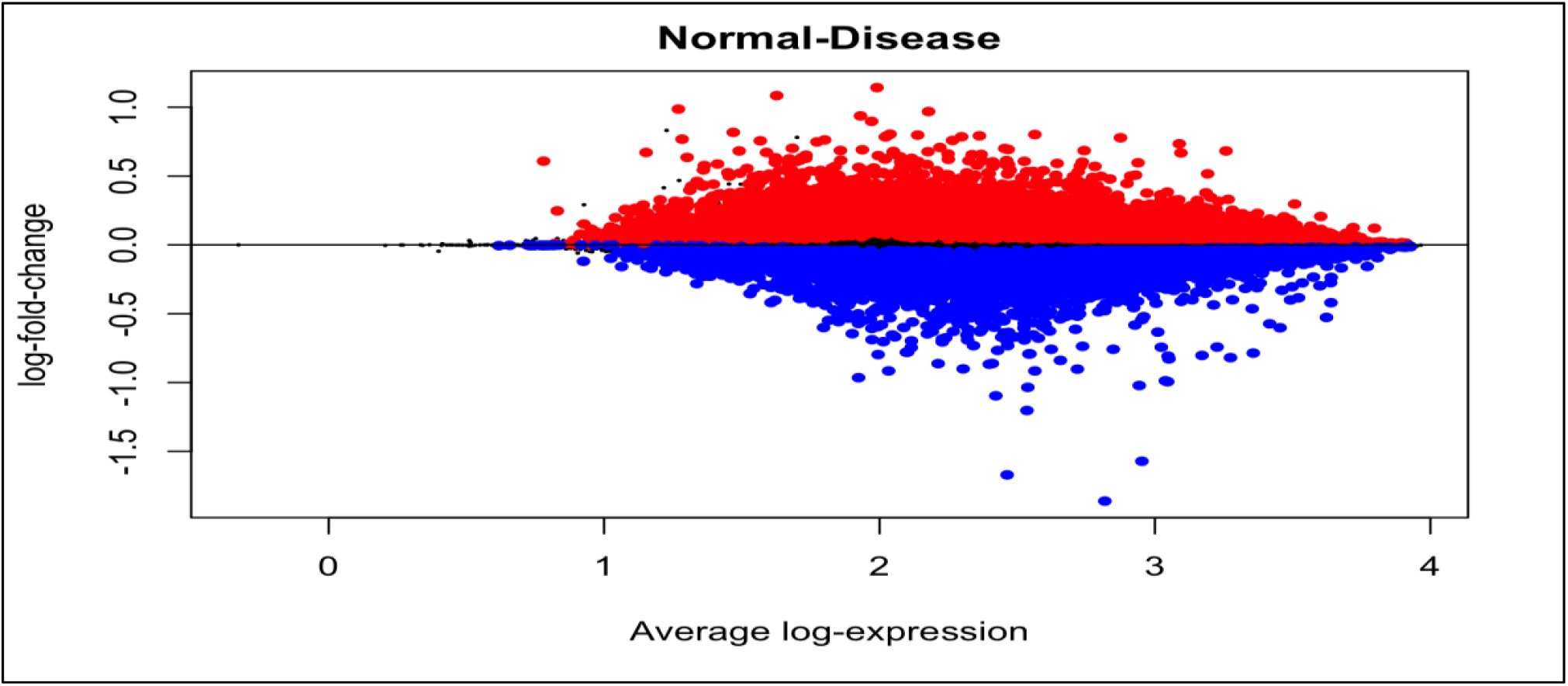
SHOWS LOGFC UPREGULATED AND DOWNREGULATED GENES

Psoriasis DEGs were combined with the NCG dataset resulting 874 up-regulated genes and 74 down-regulated genes for an in depth analysis of how differential expression can lead to malignancy. Comparing these 948 genes with cystoscope’s string melanoma disease query genes resulted in 225 genes, further these genes were filtered using Log fold change threshold of above 0.35 resulting in 18 genes which are upregulated.

### 3.2. Analysis of Topological Properties

The topological analysis of various large-scale biological networks reveals some recurring properties: power law distribution of degree, scale-freeness, and small world, which are thought to confer functional advantages such as resistance to environmental changes and tolerance to random mutations. Network analysis is an effective method for comprehending the function and evolution of biological processes, so long as smaller functional modules are equally focused on building a relationship between their topological qualities and their dynamical behaviour. The probability of degree distributions (P(K)), clustering co-efficient (C(K)), and neighbourhood connectivity (CN(K)) are topological metrics that display power law and are analysed here. The behaviour of the network is characterized by equations:

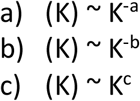

The value of the connection parameter c that is positive indicates the assortative character of the network. While the negative value of a in degree distribution indicates the availability of each network node, the positive value indicates the opposite. The -ve value of the clustering parameter b indicates dissociation in network node communication. The fundamental centrality metrics of a network, notably betweenness CB(K), closeness CC(K), and eccentricity CE(K), likewise display hierarchical behaviour.

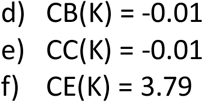

The values of the centralities exponents indicate that the network nodes have a strong regulating behaviour.

### 3.3. Network Construction and Clustering

Using stringDB mapping 28 interactions were found for 18 selected genes, degree calculation was performed using igraph R package, further genes with zero degree was removed from the list resulting in 17 genes and 28 interactions with p value of 3.13e-08.Table 1 summarises the degree for every gene. Network of Interactions were plotted using igraph R package and node size were adjusted as per degree shown in Figure 7.

**FIGURE 5.**
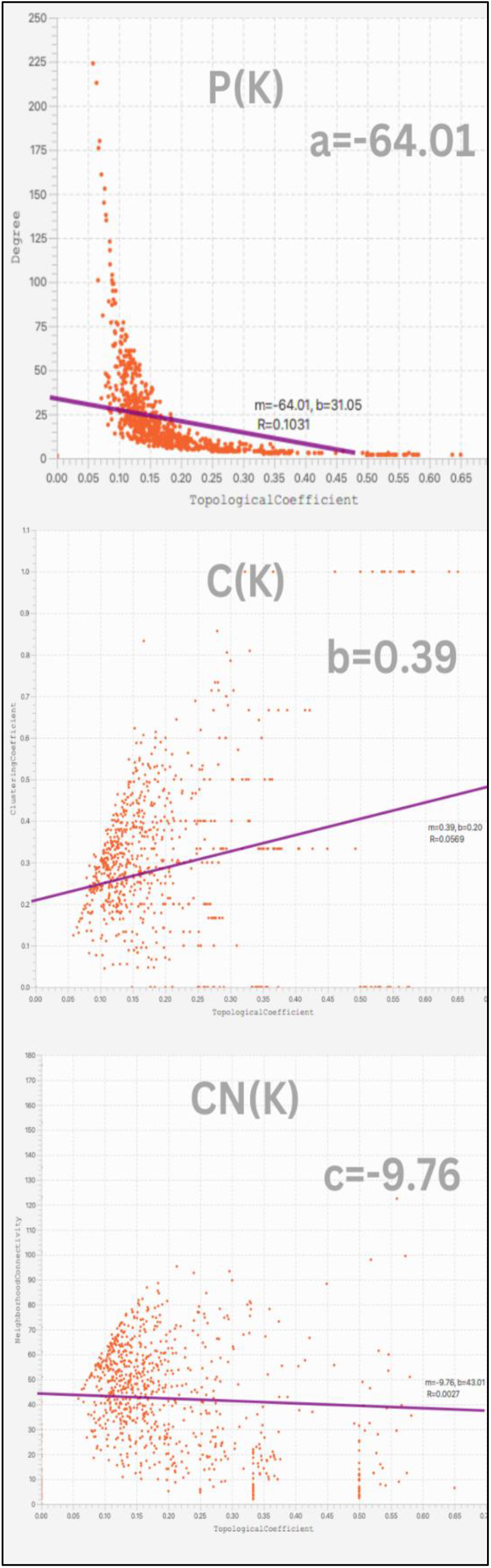
PLOTS SHOWING P(K), C(K) AND CN(K) TOPOLOGICAL PROPERTIES OF GRN.

**FIGURE 6.**
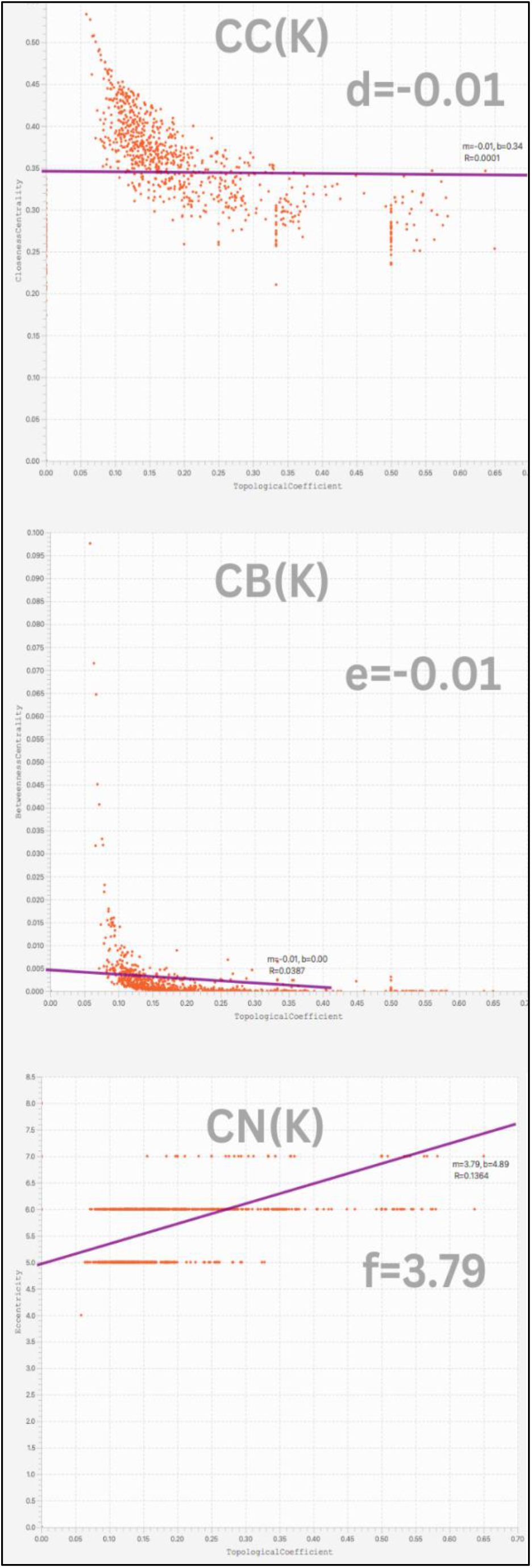
PLOTS SHOWING CC(K), CB(K) AND CE(K) TOPOLOGICAL PROPERTIES OF GRN.

**FIGURE 7.**
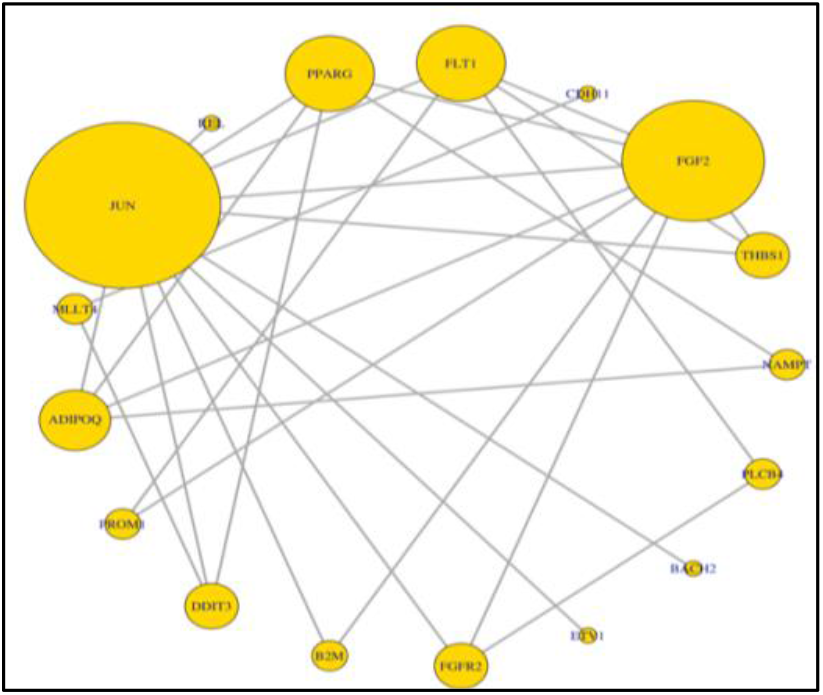
SHOWS THE INTERACTIONS OF GENES

**TABLE 1.**
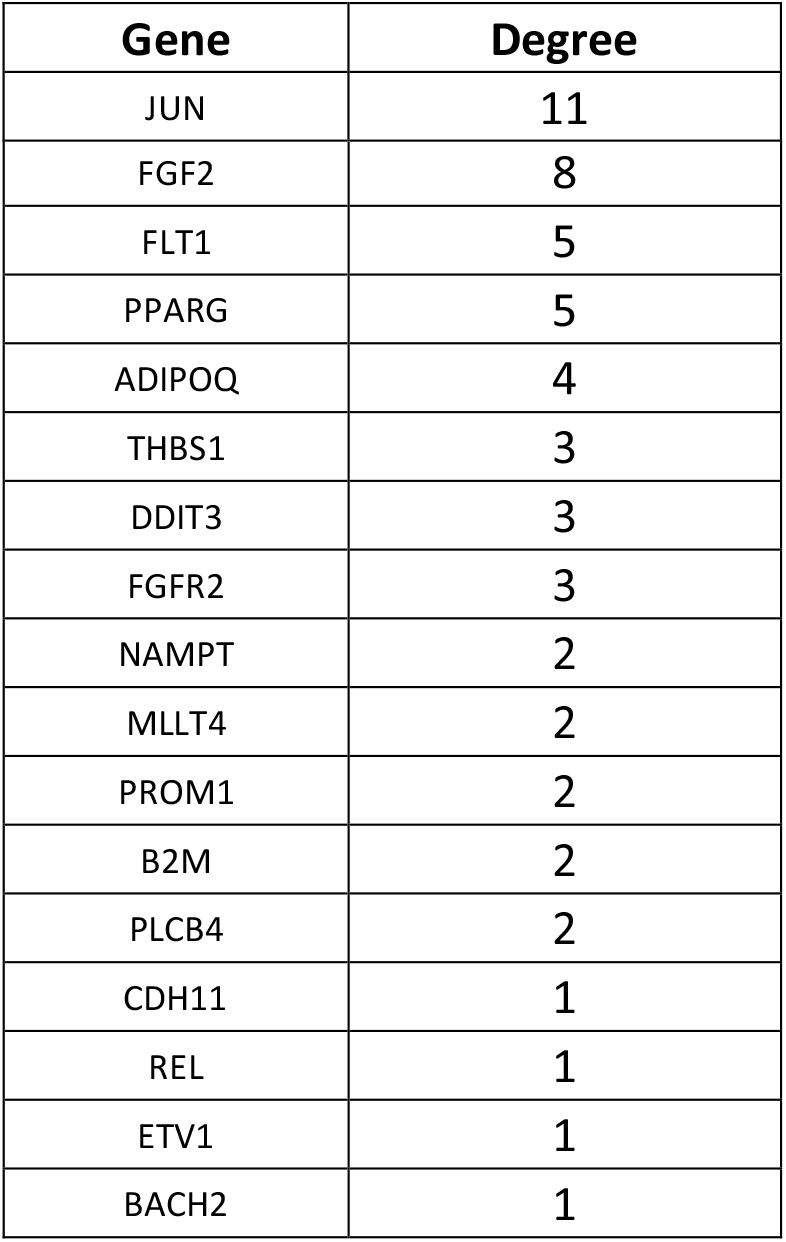
SHOWS DEGREE OF EVERY SELECTED GENE

| Gene | Degree |
| --- | --- |
| JUN | 11 |
| FGF2 | 8 |
| FLT1 | 5 |
| PPARG | 5 |
| ADIPOQ | 4 |
| THBS1 | 3 |
| DDIT3 | 3 |
| FGFR2 | 3 |
| NAMPT | 2 |
| MLLT4 | 2 |
| PROM1 | 2 |
| B2M | 2 |
| PLCB4 | 2 |
| CDH11 | 1 |
| REL | 1 |
| ETV1 | 1 |
| BACH2 | 1 |

Mutual transitivity and undirected diameter was found out to be 0.3170732 and 5 respectively. PPI enrichment score of this network calculated by strig is 6.93e-09, which is significantly more interactions than expected showing there are more interactions between proteins than would be predicted for a genome-wide random sample of proteins with the same size and degree distribution.

This enrichment shows that the proteins as a group are at least somewhat physiologically related. StringDB along with DISEASE database showed disease gene association to cancer DOID:162 with strength of 0.95, and False discovery rate of 0.0243, with 7/895 counts in network these genes are shown in red colour in Figure 8.

**FIGURE 8.**
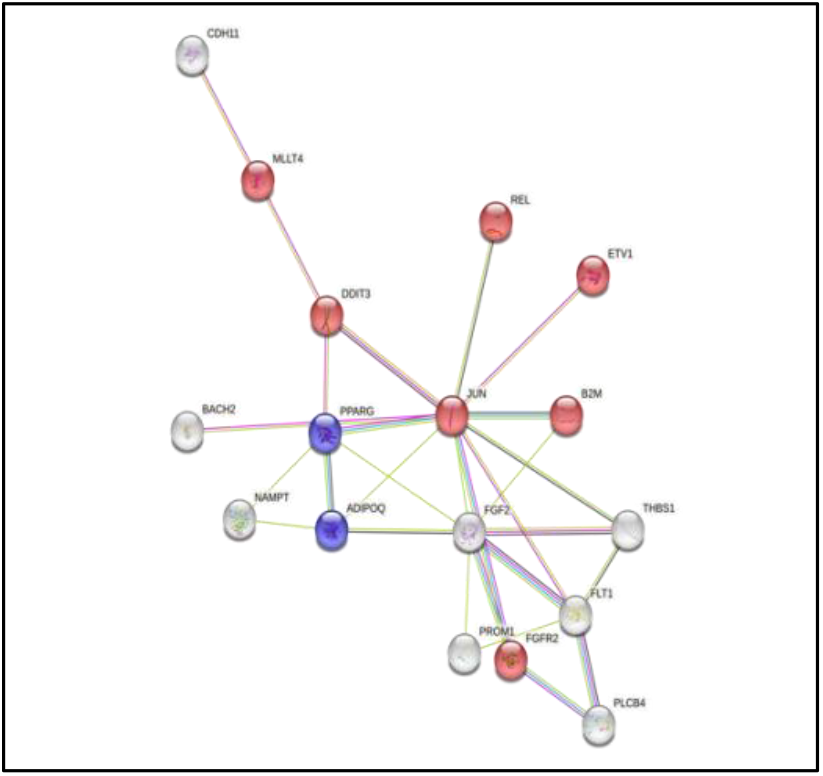
SHOWS STRINGDB INTRACTIONS WITH DISEASEDB AND TISSUEDB

StringDB along with TISSUE database showed gene expression in subcutaneous adipose tissue BTO:0004042 with strength of 2.76, and False discovery rate of 0.0202, with 2/4 counts in network these genes are shown in blue colour in Figure 8.

Igraph R package clustering tool resulted in 8 communities based on edge-between-ness with modularity of 0.1715561, dendrogram in Figure 9 shows the clustering. 4 selected Communities plotted on string network resulted in 8 hubs shown in Figure 10.

**FIGURE 9.**
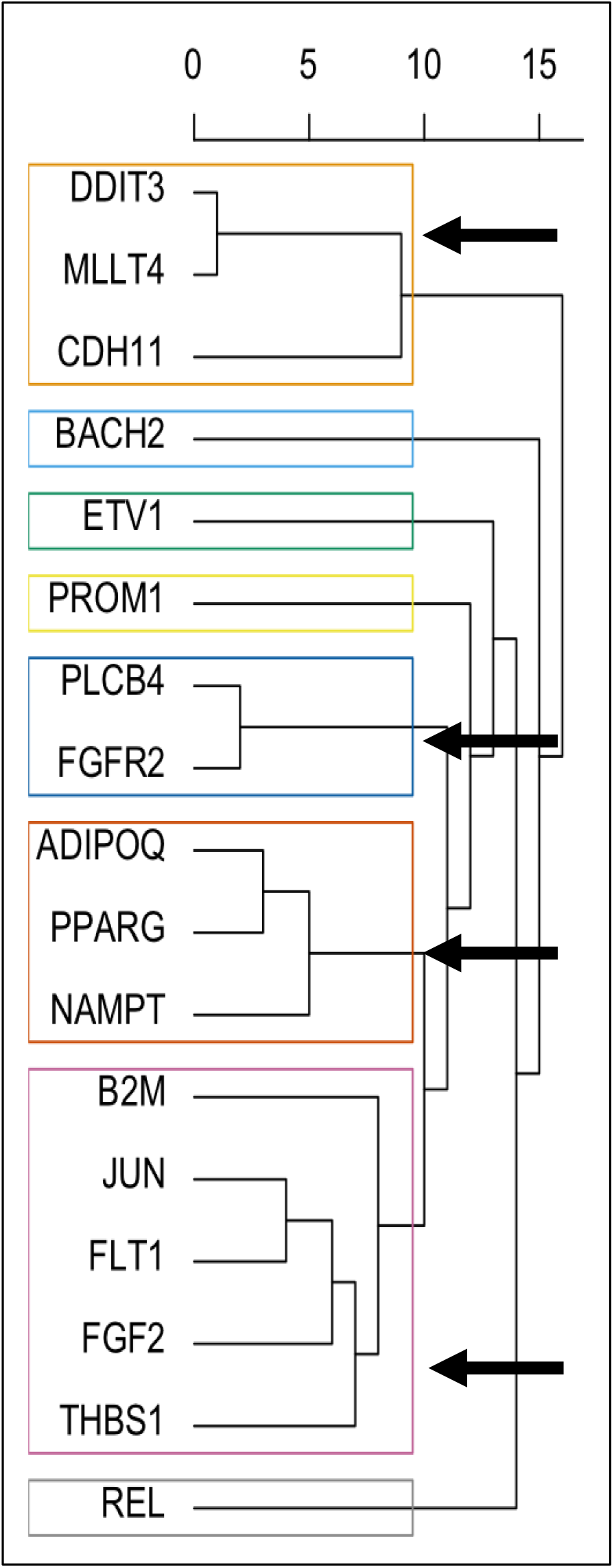
DENDROGRAM PLOT SHOWING DIFFERENT COMMUNITIES

**FIGURE 10.**
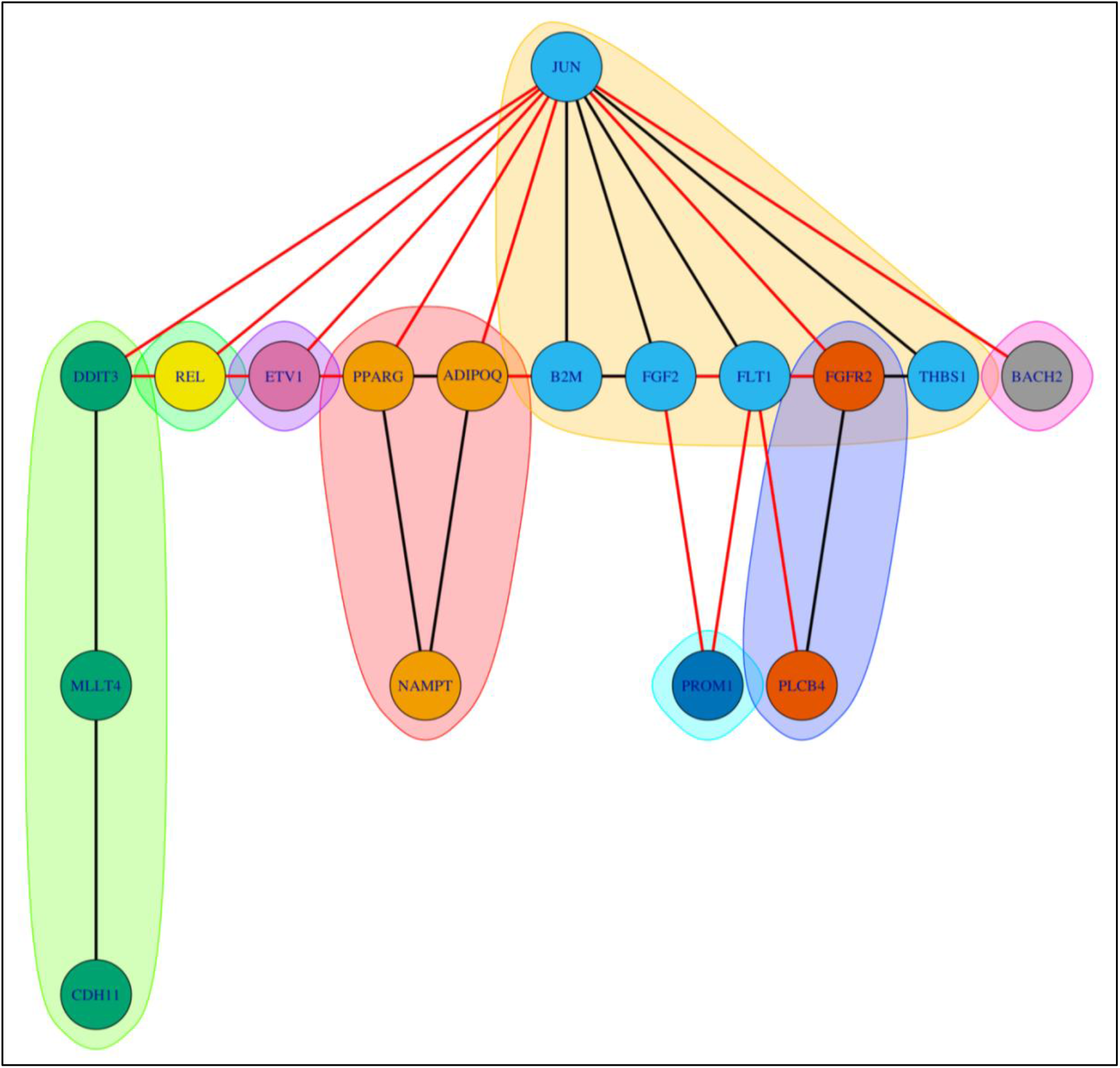
SHOWS THE COMMUNITIES ON THE STRING NETWORK

Knowing the maximum amount of interactions each node has allows us to discover possible hubs in the network. The network identified the following nodes as probable hubs: JUN, FGF2, FLT1, PPARG, ADIPOQ, THBS1, DDIT3, and FGFR2.

### 3.4. Gene Set Enrichment and KEGG pathway and Survival Analysis

To comprehend the acquired DEGs, it is necessary to understand their unique role. The microarray data-obtained genes were separated into up-regulated biomarkers (17) and down-regulated (0) elements, from which malignant genes were identified. Using cluster profiler and the Hs.org.eg.db R package, the Gene Ontology was determined for these genes, and three significant categories, namely Biological Process (bp), Cellular Component (cc), and Molecular Function (mf), were shown. It yielded results as shown in Figure 11.

**FIGURE 11.**
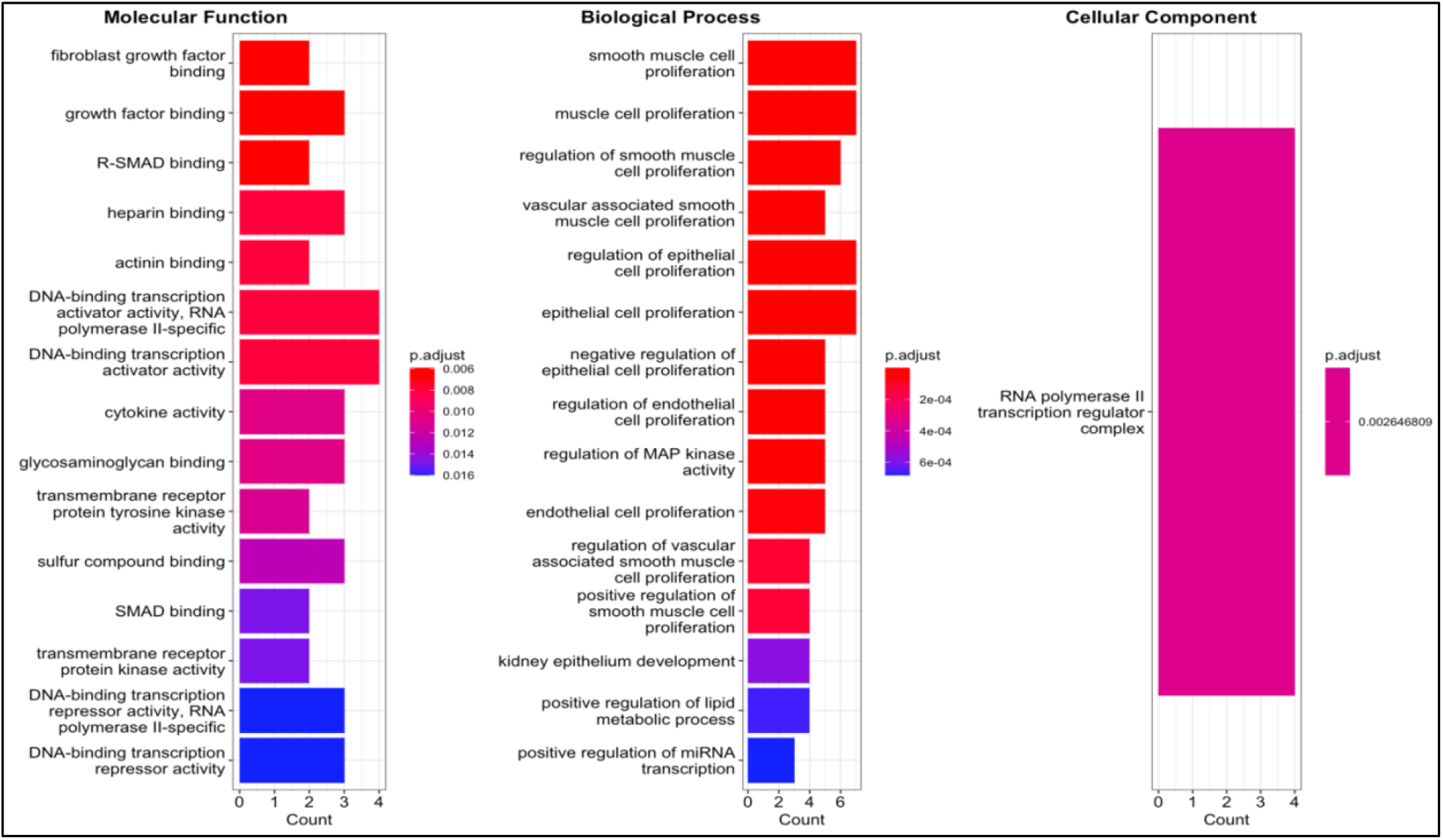
SHOWING GENE ONTOLOGY ENRICHMENT

StringDb enrichment for functional group was also performed and results are shown in Figure 12.

**FIGURE 12.**
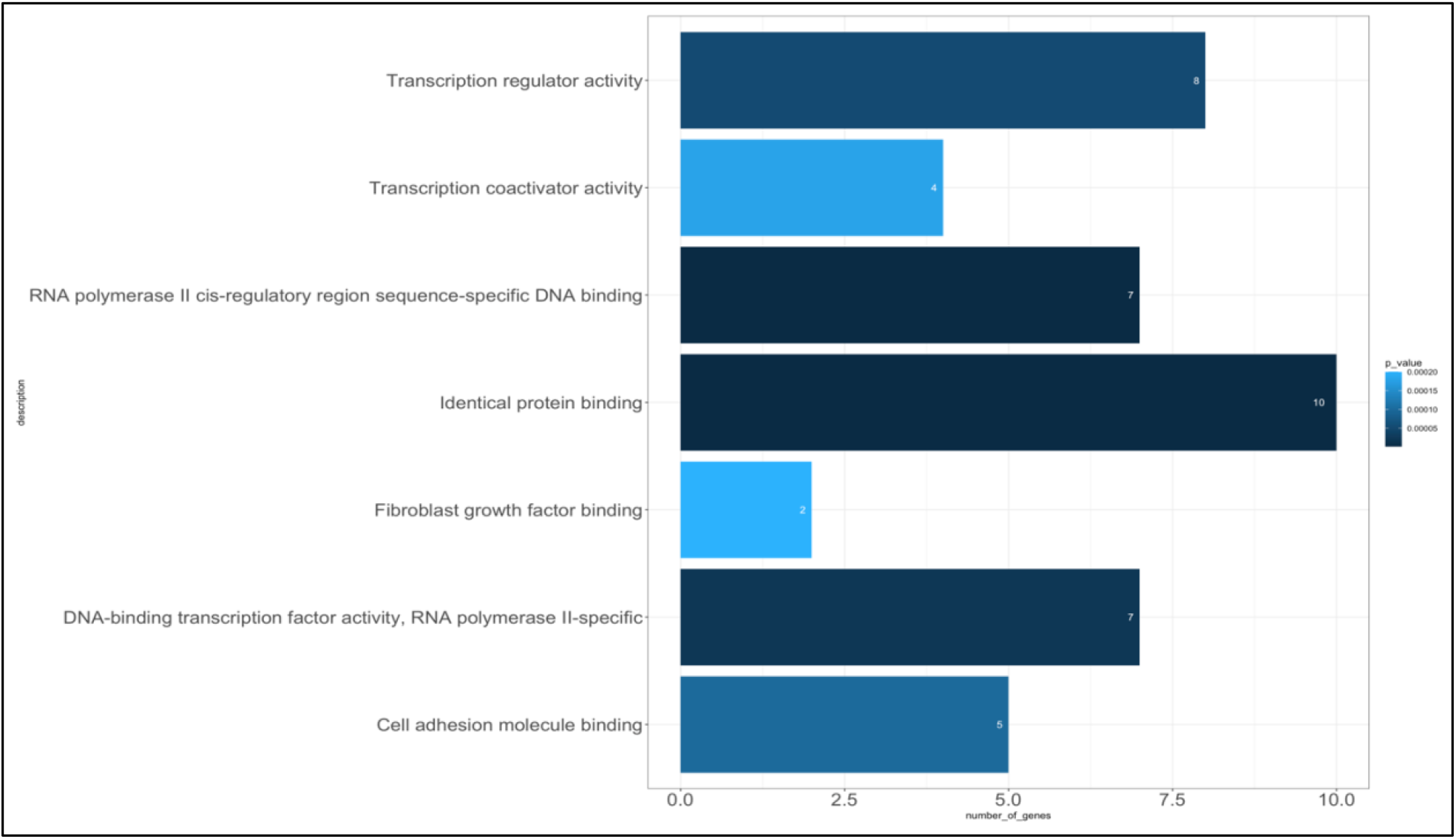
SHOWING STRINGDB FUNCTIONAL ENRICHMENT

KEGG pathway from DAVID was determined and pathways in cancer result was selected with p value 7.80e-03 and Benjamani 1.10e-01, Pathway is shown in Figure 13. Survival Analysis of Biomarker hub resulted showed that these biomarker hubs are significant in melanoma patients with patient size of 442, the data is summarised in Table 2 and survival plot is show in Figure 14. Genomic alterations are shown in Figure 15.

**Figure 13.**
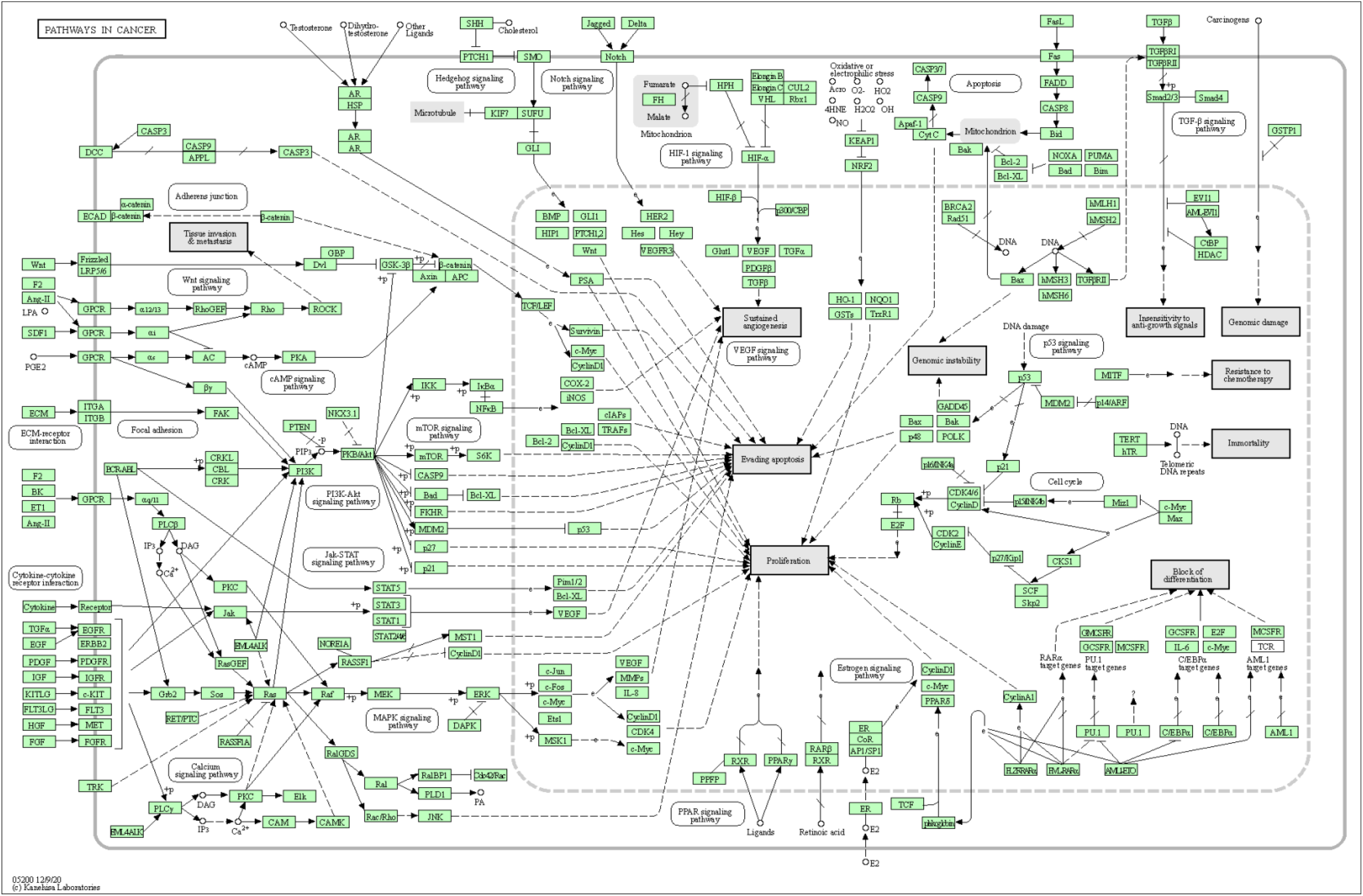
KEGG PATHWAY FOR CANCER

**FIGURE 14.**
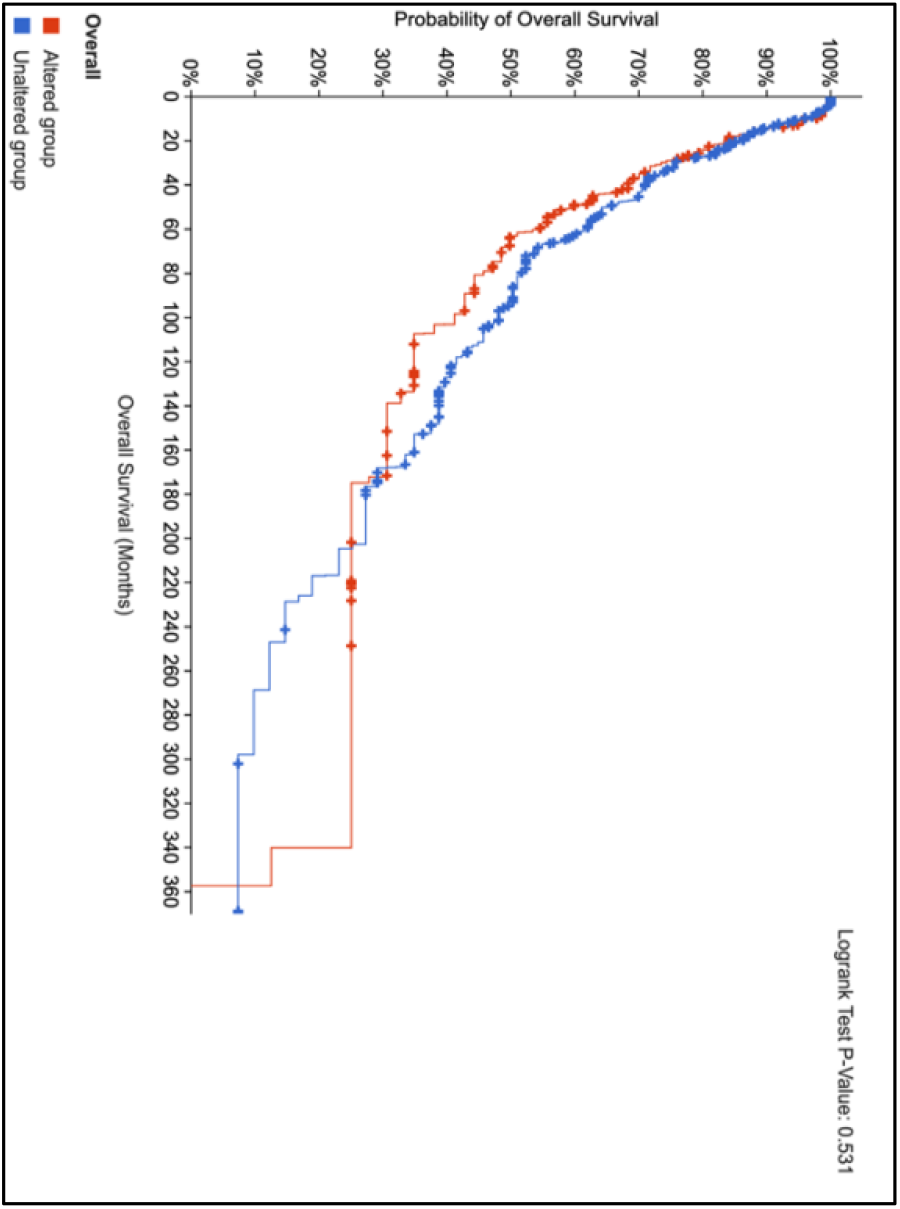
SHOWS SURVIVAL PLOT FOR 6 BIOMARKER HUB GENES

**FIGURE 15.**
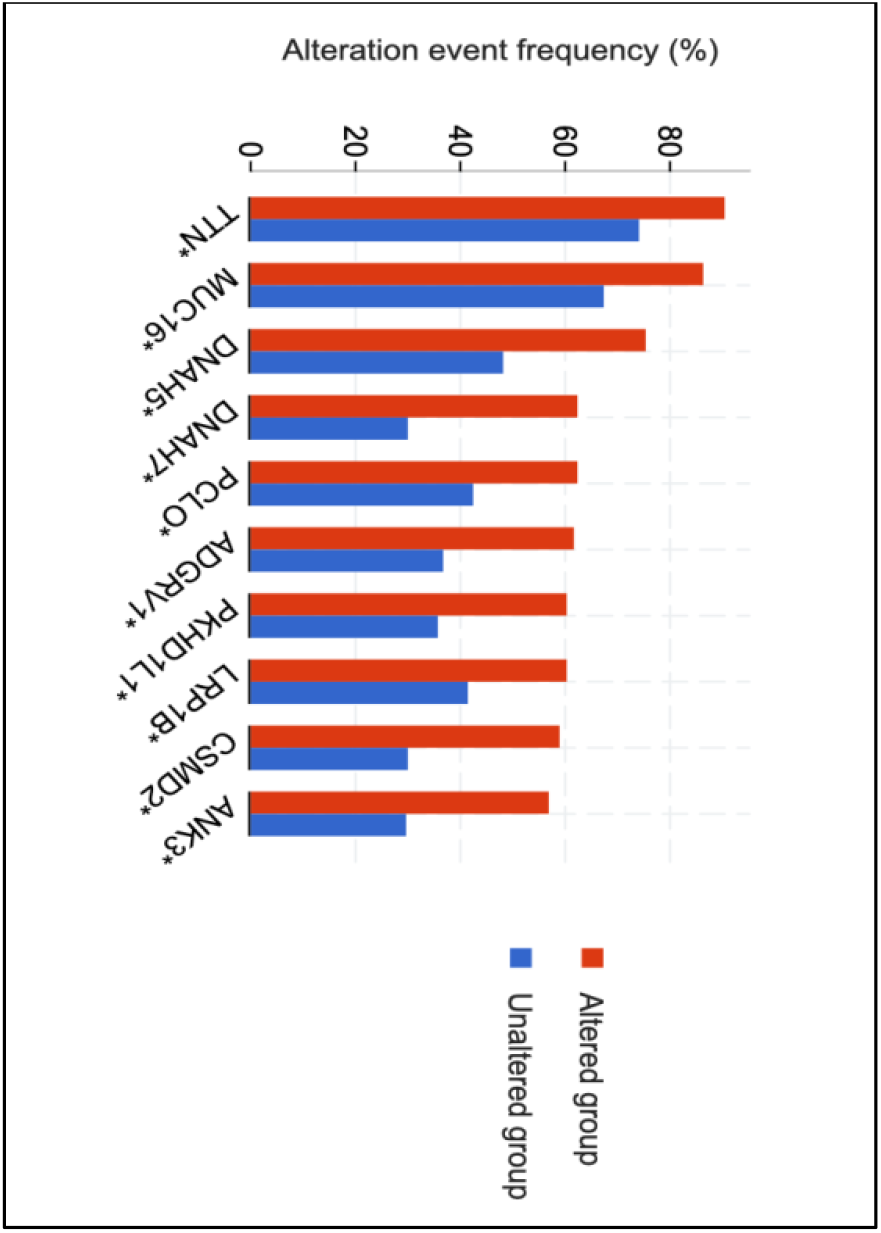
SHOWS GENE ALTERATION FOR 6 BIOMARKER HUB GENES

**Table 2.** summarises survival analysis for 6 biomarker hubs in 442 patients

| Survival Type | Progression Free | Overall | Disease-specific | Disease Free |
| --- | --- | --- | --- | --- |
| Number of Patients | 427 | 426 | 420 | 0 |
| # in Altered group | 139 | 139 | 136 | 0 |
| # in Unaltered group | 288 | 287 | 284 | 0 |
| Median months survival in Altered group (95% CI) | 32.84 (21.80 - 43.82) | 63.02 (50.89 - 103.26) | **80.68 (58.52 - 138.80)** | NA |
| Median months survival in Unaltered group (95% CI) | 37.87 (31.07 - 47.37) | 93.01 (66.67 - 117.93) | **105.04 (69.07 - 152.81)** | NA |
| p-Value | 0.168 | 0.531 | 0.605 | N/A |
| q-Value | 0.503 | 0.605 | 0.605 | N/A |

## 4. CONCLUSIONS

This research attempts to provide light on the regulatory network of psoriasis and melanoma cancer. The communities (and sub-networks) give insight into the relationships between the biomarkers that contribute to the development of cancer. It can be shown that possible biomarkers for melanoma, such as JUN, FGF2, FLT1, PPARG, ADIPOQ, THBS1, DDIT3, and FGFR2, are present in the network up to the higher levels. Also, the topological features imply two things: degree distribution plots (P(K), C(K), CN(K)) imply the presence of hierarchy in the network, whereas centralities (CB(K), CC(K), CE(K)) imply the assortative nature of the network. The existence of biomarkers in higher levels, the plot for modularity and the topological qualities combined show the presence of possible hubs at every level of the hierarchy of network. The relevance of survival study indicates that these biomarkers play a vital role in melanoma. In conclusion, this study may be utilised for future research to identify prospective therapeutic targets and, when further developed, can lead to the creation of more effective treatments for melanoma. Also This study sheds light on the relationship between psoriasis and melanoma malignancy.

## 6. Acknowledgement

We would like to express our gratitude towards Department of Oncology, Shree Aggarsain International Hospital for their support and I would like to thank Dr. S. Kumar and Dr. V. Aggarwal, for guiding me in this research.

## 8. Data Availability

Primary Dataset is available on NCBI’s GEO (Gene expression Omnibus, http://www.ncbi.nlm.nih.gov/geo/GSE14905), all the other generated data is avaible with primary author.

